# Immunogenetic response to a malaria-like parasite in a wild primate

**DOI:** 10.1101/2020.11.24.396317

**Authors:** Amber E. Trujillo, Christina M. Bergey

**Affiliations:** Department of Anthropology, New York University, 25 Waverly Place, New York, NY, 10003; New York Consortium in Evolutionary Primatology; Department of Genetics, Rutgers University, 604 Allison Road, Piscataway, NJ, 08840

## Abstract

Malaria is infamous for the massive toll it exacts on human life and health. In the face of this intense selection, many human populations have evolved mechanisms that confer some resistance to the disease, such as sickle-cell hemoglobin or the Duffy null blood group. Less understood are adaptations in other vertebrate hosts, many of which have a longer history of co-evolution with malaria parasites. By comparing malaria resistance adaptations across host species, we can gain fundamental insight into host-parasite co-evolution. In particular, understanding the mechanisms by which non-human primate immune systems combat malaria may be fruitful in uncovering transferable therapeutic targets for humans. However, most research on primate response to malaria has focused on a single or few loci, typically in experimentally-infected captive primates. Here, we report the first transcriptomic study of a wild primate response to a malaria-like parasite, investigating gene expression response of red colobus monkeys (*Piliocolobus tephrosceles*) to natural infection with the malaria-like parasite, *Hepatocystis*. We identified colobus genes with expression strongly correlated with parasitemia, including many implicated in human malaria and suggestive of common genetic architecture of disease response. For instance, the expression of *ACKR1* (alias *DARC*) gene, previously linked to resistance in humans, was found to be positively correlated with parasitemia. Other similarities to human parasite response include induction of changes in immune cell type composition and, potentially, increased extramedullary hematopoiesis and altered biosynthesis of neutral lipids. Our results illustrate the utility of comparative immunogenetic investigation of malaria response in primates. Such inter-specific comparisons of transcriptional response to pathogens afford a unique opportunity to compare and contrast the adaptive genetic architecture of disease resistance, which may lead to the identification of novel intervention targets to improve human health.

**Author Summary:** The co-evolutionary arms race between humans and malaria parasites has been ongoing for millennia. Fully understanding the evolved human response to malaria is impossible without comparative study of parasites in our non-human primate relatives. Though laboratory primates are fruitful models, the complexity of wild primates infected in a natural transmission system may be a more suitable comparison for contextualizing malaria infections in human patients. Here, we investigate the genetic mechanisms underlying the immune response to *Hepatocystis*, a close relative of human-infective malaria, in a population of wild Ugandan red colobus monkeys. We find that the genes involved have considerable overlap with those active in human malaria patients. Like *Plasmodium*, *Hepatocystis* induces changes in blood cell type and may cause the host to produce blood components outside of the bone marrow or alter metabolism related to the production of lipids. Our work helps to identify the genetic mechanisms underlying the arms race between primates and malaria parasites, providing fundamental evolutionary insight. Such comparative work on the interaction between wild non-human primates and malaria parasites can identify ways in which primates have evolved resistance to malaria parasites, and further investigation of such implicated genes may lead to novel potential therapeutic and vaccine targets.

## Introduction

Though malaria parasites (order Haemosporida) infect a broad variety of vertebrates via a broad variety of dipteran insect vectors (Galen et al., 2018; Perkins, 2014), the five species that have caused incalculable human deaths have understandably garnered the most research attention. Evidence of this lengthy battle is evident from scars in the human genome, as malaria has exerted one of the strongest selection pressures on the human immune system (Kwiatkowski, 2005; Kariuki and Williams, 2020). Recent investigations simultaneously assaying human and malaria gene expression in the blood have illuminated the genetic architecture of this host-parasite interaction, permitting fundamental insight into the immunology of this disease (Yamagishi et al., 2014; Otto et al., 2018; Lee et al., 2018; Mukherjee and Heitlinger, 2019). However, a full understanding of the adaptive human immunogenetic response to malaria requires comparative data from closely related primates.

Few investigations of malaria and malaria-like parasite response in non-human primates have been undertaken, and fewer still have investigated gene expression response in free-ranging individuals in a natural pathogen transmission system. Most studies in non-human primates have instead made use of experimentally-infected laboratory primates (*e.g.*, rhesus macaques, *Macaca mulatta*; Ylostalo et al., 2005; Tang et al., 2017, 2018; Galinski, 2019; Tang et al., 2019). However, tightly controlled laboratory infections likely do not capture the complexity in parasite behavior and host response characterizing natural infections, as has been found for human patients (Daily et al., 2007). In one of the few studies of malaria in wild primates, variation in the *ACKR1* (alias *DARC*) gene was found to be associated with phenotypic diversity in susceptibility to *Hepatocystis* (a malaria-like parasite) in yellow baboons (*Papio cynocephalus*; Tung et al., 2009). High throughput transcriptomic profiling of host-parasite interactions in wild primates living in natural transmission systems is needed to fully understand the evolution of malaria response in primates.

Contained within the polyphyletic *Plasmodium* clade (Galen et al., 2018), members of genus *Hepatocystis* commonly parasitize bats and non-human primates. *Hepatocystis* is often highly prevalent in primates (20% to 90% across host species; Thurber et al., 2013; Ayouba et al., 2012), and parasitemia may persist for over a year without clinical symptoms (Vickerman, 2005) though reports exist of anemia and liver scarring due to monocytes infiltration (Garnham, 1966). At least one species is able to host-switch between primate species (Thurber et al., 2013), but no human infections have been reported. Though nested within the *Plasmodium* genus as a sister taxon to bat- and rodent-infecting *Plasmodium* in particular (Galen et al., 2018), *Hepatocystis* parasites differ in their apparently lack of asexual replication in red blood cells and their use of midges as vectors (Aunin et al., 2020; Garnham, 1966).

Here, we report the first study of genome-wide gene expression response to a malaria-like parasite in wild non-human primates. We investigate the genetic architecture of response to *Hepatocystis* in a population of wild red colobus monkeys (*Piliocolobus tephrosceles*) living in Kibale National Park, Uganda. We identify colobus monkey and *Hepatocystis* genes and genetic pathways that show differential expression in response to *Hepatocystis* infection. We find considerable overlap between the genes involved in the colobus response to *Hepatocystis* and human response to *Plasmodium* malaria. Overall, our study illustrates the importance of comparative host-parasite genomic investigations for understanding the genetic architecture of human disease response.

## Results and Discussion

The Ugandan red colobus monkey whole blood transcriptomic dataset was collected by Simons and colleagues (2019) as part of the effort to generate the red colobus monkey reference genome. The sampled individuals (*N* = 29) are from a habituated population in Kibale National Park, Uganda, where red colobus have been the subjects of many disease-related studies as a part of the the Kibale EcoHealth Project (Chapman et al., 2005, 2013; Goldberg et al., 2008; Lauck et al., 2011; Simons et al., 2019; Aunin et al., 2020). To identify colobus monkey genes with expression correlated to *Hepatocystis* parasite levels, we aligned blood transcriptome reads to a concatenated host-parasite pseudo-reference genome (containing genomes of red colobus and *Hepatocystis* sp.) and tallied reads by gene. After filtering, the median per-sample count of reads uniquely mapping to the colobus genome was 9,531,580 (range: 4,149,243 to 12,907,207). We next computed a proxy of *Hepatocystis* parasitemia as the ratio of reads mapped uniquely to the primate versus parasite genome (Fig. 1; Table S1). For *Hepatocystis* reads, we used published read counts for these samples (Aunin et al., 2020) excluding genes with expression in the top and bottom quartile to minimize the impact of genes with outlier expression.

**Figure 1.**
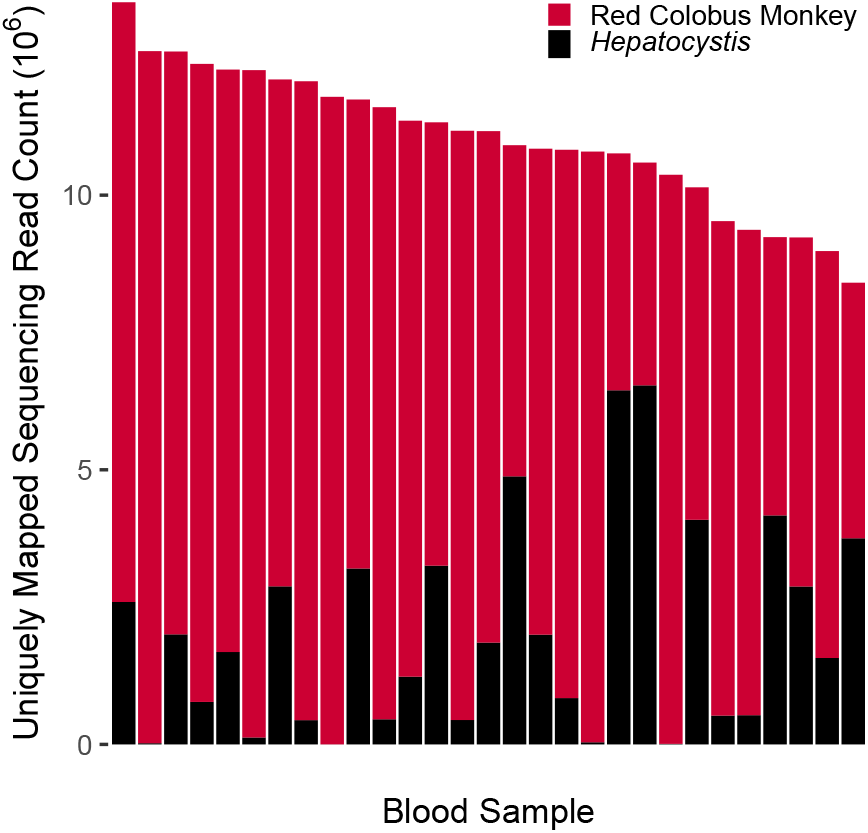
Uniquely mapped read counts from red colobus monkey (red) and *Hepatocystis* parasite (black) in monkey blood samples. *Hepatocystis* counts based on those of Aunin et al., 2020, excluding genes with expression in the highest and lowest quartiles. The ratio of *Hepatocystis* to colobus reads for each individual was used as a proxy for parasitemia.

### *Hepatocystis* parasitemia correlates with proportions of monocytes and neutrophils

To investigate the cellular composition of the red colobus immune response, we first tested if the relative abundance of red colobus monkey immune cell type subsets (Simons et al., 2019) correlated with inferred *Hepatocystis* parasitemia (Table S2; Fig. 2, S1). Though *Hepatocystis* differs from some *Plasmodium* malaria species in causing long-lasting, chronic infections (Garnham, 1966), our findings suggest *Hepatocystis* parasite load causes similar changes in blood cell counts. In *Plasmodium* malaria, such changes in hematologic parameters have been hypothesized to be associated with changes in the host immunity or ability to mount an effective immune response (Ingersoll et al., 2010).

**Figure 2.**
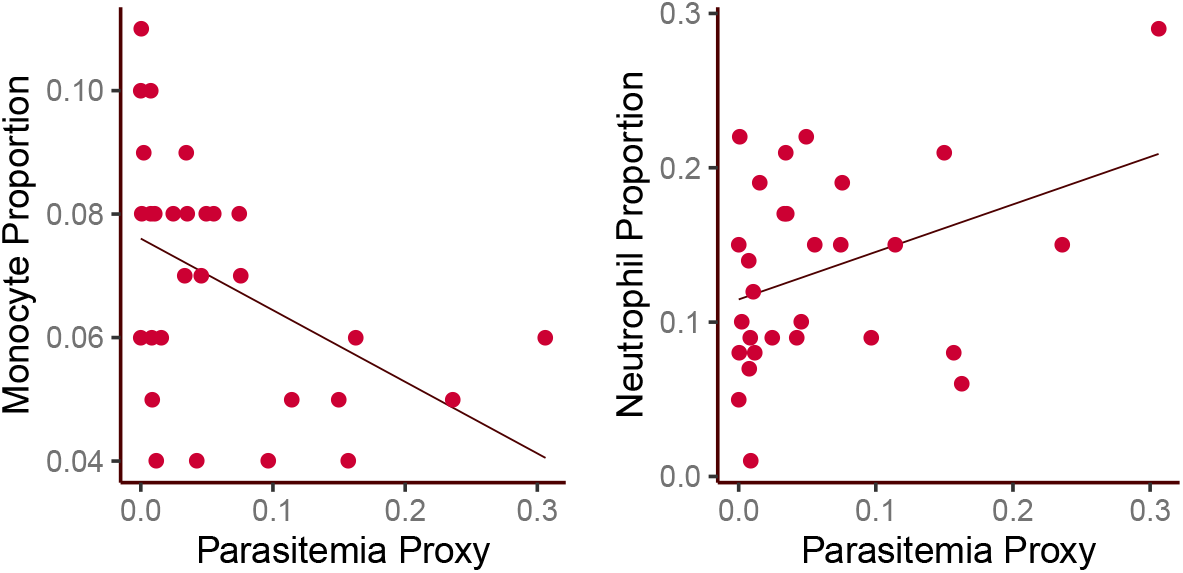
Monocyte and neutrophil proportions significantly associated with parasitemia.. Relationship between proportions of (left) monocytes and (right) neutrophils with inferred *Hepatocystis* parasitemia. The *p*-values listed summarize *t*-test for significant association (*e.g.*, Pearson’s correlation coefficient not equal to zero). Proportions of monocytes and neutrophils were significantly negatively and positively correlated with parasite load, respectively.

The relative proportion of monocytes was significantly negatively associated with parasitemia (*p* = 0.0134, Pearson’s product moment correlation coefficient *r*^2^ = −0.454; Fig. 2). Unsurprisingly, given their involvement in protection from diverse pathogens, monocytes have central roles in the innate immune response to malaria, with involvement in diverse processes including phagocytosis, cytokine production and antigen presentation (Ortega-Pajares and Rogerson, 2018). However, products of *Plasmodium*, including hemozoin and extracellular vesicles, may impair monocyte function. Investigations of the relationship between malaria parasitemia and monocyte proportion in humans have yielded contrasting results with some studies reporting low monocyte counts in relation to high parasitemia (Kotepui et al., 2015) similar to our findings, while others report the opposite (Berens-Riha et al., 2014). This apparent contradiction may be due to differences in the hematological profile of individuals from different locales (Gansane et al., 2013).

In contrast to monocytes, the proportion of neutrophils was significantly positively correlated with parasitemia (*p* = 0.0469, *r*^2^ = 0.372; Fig. 2). Differences in observed response to malaria parasitemia have also been observed in neutrophil counts for cohorts from different geographic areas, possibly attributable to immune status or age (Aitken et al., 2018). Our findings are similar to those of a comparative study of uncomplicated malaria in pediatric patients across sub-Saharan Africa, in which neutrophil counts were found to be significantly positively correlated with parasitemia (Olliaro et al., 2011).

Similarly, we next tested whether the proportion of *Hepatocystis* parasites in particular stages of the parasite life cycle was correlated with parasite load. Using the recently generated single cell transcriptomic data for *Plasmodium berghei* malaria (Howick et al., 2019), we deconvoluted the bulk *Hepatocystis* gene expression data in order to infer parasite life stage proportions for each sample (after Aunin et al., 2020; Table S3; Fig. S2). We found no significant correlation between inferred parasitemia and parasite life stage proportion (Table S4; Fig. S3). This is consistent with expectations for chronic *Hepatocystis* infection, in which parasite life stages persist throughout the infection, potentially due to constant generation of new parasites rather than re-infection (Aunin et al., 2020).

Finally, we tested whether relative abundances of *Hepatocystis* developmental stages and colobus immune cell types were correlated. The proportion of memory B cells was significantly negatively correlated with *Hepatocystis* female gametocytes (*r*^2^ = −0.416, *p* = 0.0246; Table S5). Our results are consistent with findings that memory B cells dominate early response after a malaria rechallenge in mice, while sexual stages of the lifecycle do not appear until late infection (Krishnamurty et al., 2016).

### Numerous genes are implicated in immunogenetic response to *Hepatocystis*

Of the 12,223 red colobus genes considered after filtration, 324 and 235 genes were significantly positively or negatively correlated with inferred parasitemia, respectively, allowing a False Discovery Rate (FDR) of 20% (Fig. 3). The red colobus genes with expression most strongly positively correlated with parasitemia (Table 1, S6; Fig. 4) included *UBE2K*, *PP2D1*, *TMEM167A*, *LSM14A*, *UBE2B*, *TTLL12*, *AGFG1*, and *APOBEC2*.

**Table 1.**
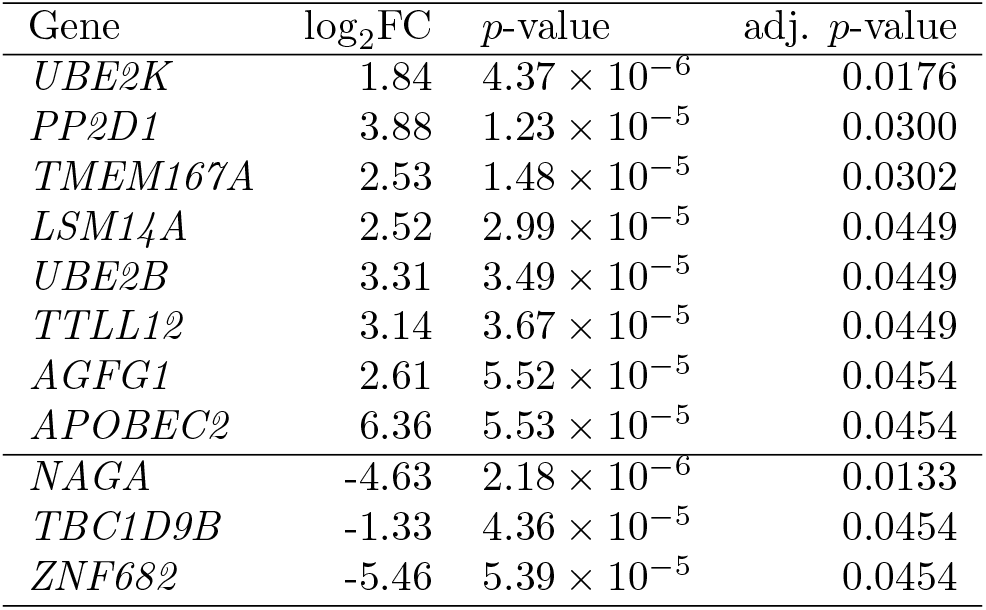
Red colobus monkey genes with expression most strongly correlated with parasitemia. The log_2_ fold change in response to inferred parasitemia (“log_2_FC”) is computed per unit change in inferred parasitemia proxy. All genes lacking orthology information (*i.e.*, with identifiers beginning with “LOC”) were excluded from this table. Genes with Benjamini-Hochberg FDR-adjusted *p*-value *<* 0.05 show. Full results available in Table S6.

**Figure 3.**
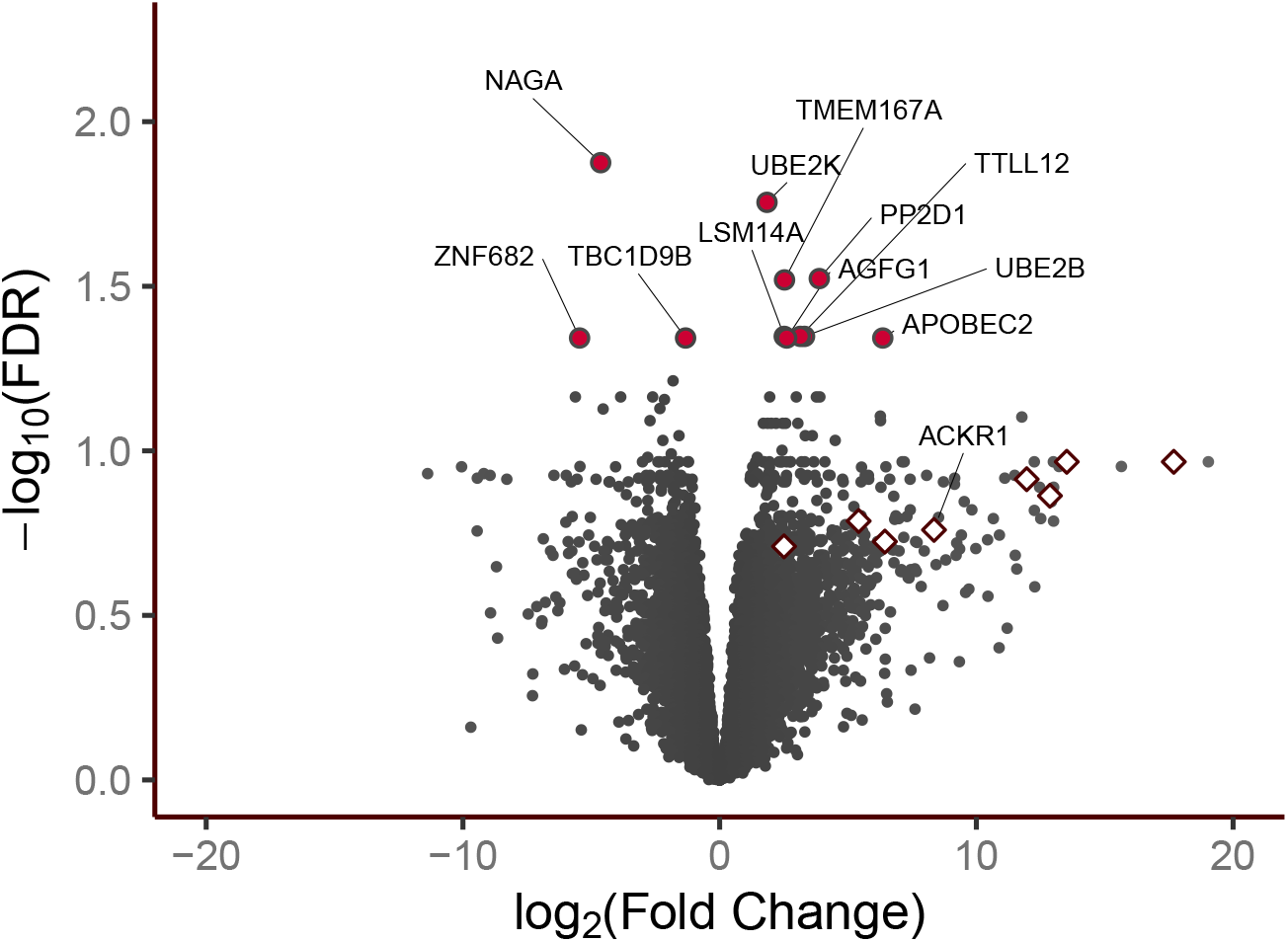
Volcano plot showing magnitude and significance of colobus gene expression changes correlated with inferred *Hepatocystis* parasitemia. Labelled red circles indicate colobus genes significantly up- or down-regulated in response to parasitemia (*p <* 0.05 after Benjamini-Hochberg adjustment for FDR). The log_2_ fold change in response to inferred parasitemia (x-axis) is computed per unit change in inferred parasitemia proxy. Diamond points indicate genes associated with erythrocyte-associated disorders such as stomatocytosis that were enriched among up-regulated genes. All genes lacking orthology information (*i.e.*, with identifiers beginning with “LOC”) were excluded.

**Figure 4.**
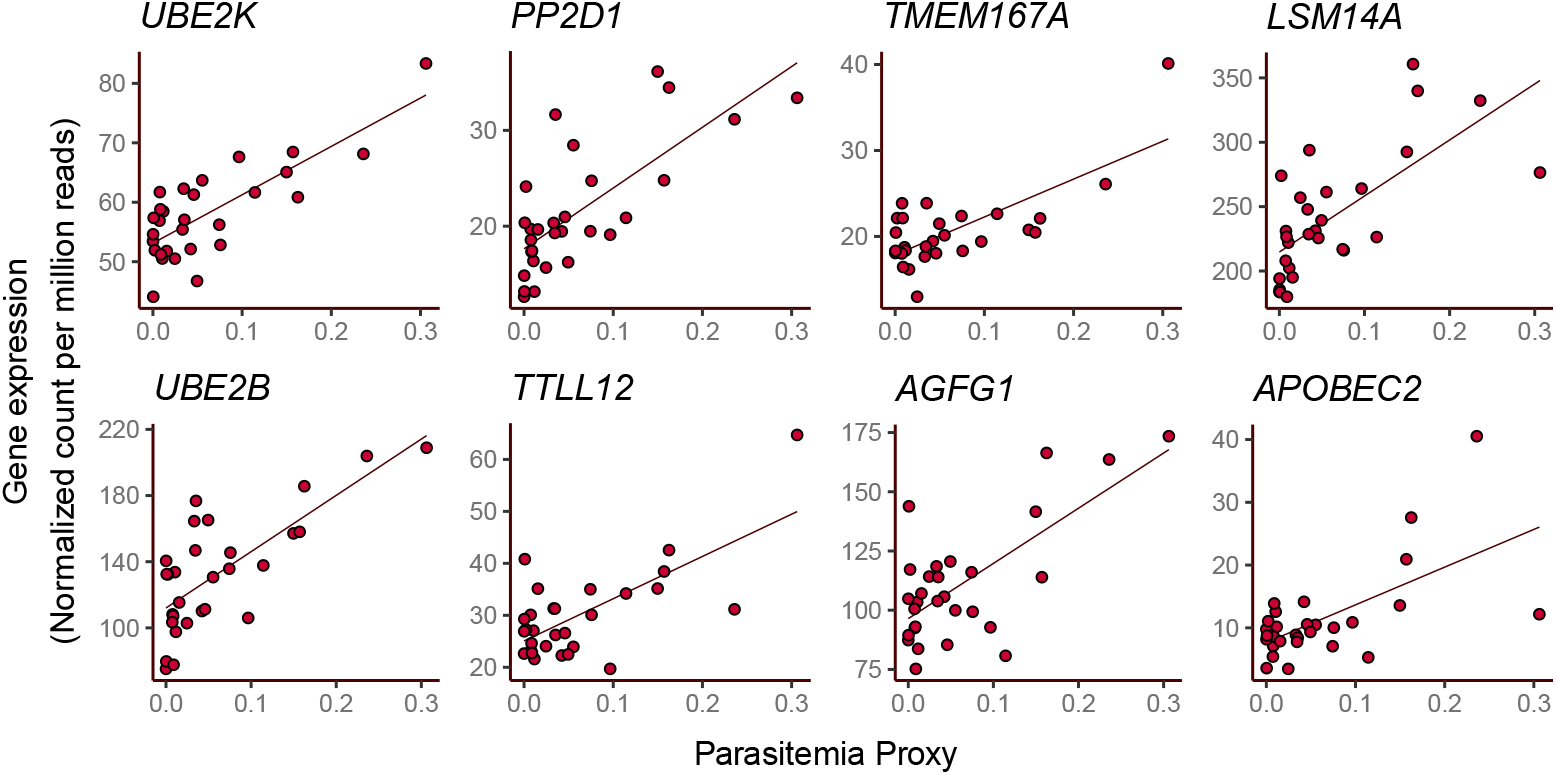
Expression of significantly up-regulated colobus genes positively correlated with *Hepatocystis* parasitemia. Expression summarized as normalized read count per million reads sequenced.

These genes with the strongest evidence of being up-regulated in response to *Hepatocystis* parasitemia were often associated with red blood cell phenotypes or malaria response. For instance, *LSM14A* has been linked to red blood cell (RBC) distribution width (*i.e.*, red blood cell volume variation; Kichaev et al., 2019) and antiviral cytokine induction (Liu et al., 2016), and likely contributes to stress granule formation in mice infected with *Plasmodium* (Hanson and Mair, 2014). *APOBEC2* is a part of the APOBEC family of (deoxy)cytidine deaminases which play an integral role in innate immunity (Jha et al., 2012) and were found to be upregulated in early liver-stage *Plasmodium* infection in mice (Albuquerque et al., 2009).

The red colobus genes with expression most strongly negatively correlated with parasitemia (Table 1, S6; Fig. 5) included *NAGA*, *TBC1D9B*, and *ZNF682*.

**Figure 5.**
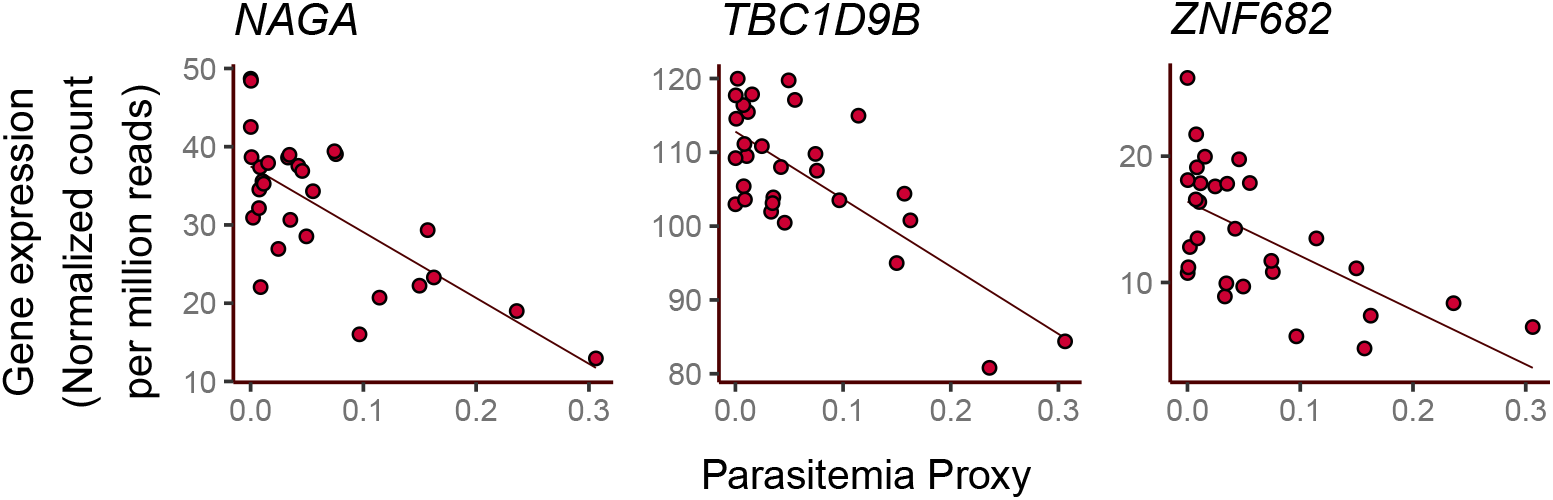
Expression of significantly down-regulated colobus genes negatively correlated with *Hepatocystis* parasitemia. Expression summarized as normalized read count per million reads sequenced.

We were particularly interested in the relationship between *Hepatocystis* parasitemia and expression of Atypical Chemokine Receptor 1 (*ACKR1*, alias *DARC*) which defines the Duffy blood group. *ACKR1* is the only gene associated with *Hepatocystis* protection in a wild primate population, with polymorphisms in its promoter region associated with variation in susceptibility to *Hepatocystis kochi* in yellow baboons (*Papio cynocephalus*; Tung et al., 2009). In the red colobus monkey population, we found that the *ACKR1* (*DARC*) gene expression was positively correlated with parasitemia (*p* = 0.00351, adjusted *p* = 0.159; Fig. S4). The Duffy antigen is an important receptor by which *Plasmodium vivax* and *P. knowlesi* infect human and non-human primate erythrocytes, and humans with the Duffy negative blood group—meaning cells that do not express this receptor—are semi-protected from infection by these malaria species (Miller et al., 1975; Mason et al., 1977; Michon et al., 2001). Underscoring the selective advantage and evolutionary importance of this gene in humans, the allele that confers protection against malaria (a variant at a single polymorphic site in the promoter region; Tournamille et al., 1995) has independently arisen in humans in malaria endemic regions, particularly Papua New Guinea and parts of sub-Saharan Africa (Miller et al., 1975; Zimmerman et al., 1999), and has rapidly reached high frequency following movement of people into areas with high *P. vivax* prevalence (Pierron et al., 2018; Hodgson et al., 2014). Our results add to the prior evidence (Tung et al., 2009) that the alleles at the *ACKR1* gene may confer a selective advantage to non-human primates mediated by *Hepatocystis* susceptibility, similar as has been established in humans for malaria-mediated selection.

Strategies for host organism response to infection and mitigation of pathology include alteration of immune cell type proportions. To investigate whether the differential expression we observed in response to parasitemia was wholly or in part due to changes in cell-type composition, we added all relative cell-type proportions alongside the parasitemia proxy in the model of read counts. We found the results of the test for differentially expressed genes were broadly similar to the simple model (Table S7; Fig. S5). Very few genes that had been associated with parasitemia in the simple model were no longer significant when relative immune cell-type proportions were included as parameters, which would indicate their response to parasitemia may be indirect and associated with changes in cell-type composition. Of the top 12 named genes associated with the parasitemia proxy with the highest confidence (uncorrected *p <* 0.0001), only two (*LSM14A* and *NAGA*) remained associated at that strict significance cutoff, though under a more lenient cutoff of *α* = 0.01 to allow for the decrease in statistical power associated with the more complex model, 10 of the 12 original genes were significant (*TTLL12* and *ZNF682* were not). Our results suggest that genes such as *LSM14A* and *NAGA* are cell composition-independent in their response to parasitemia, but other genes such as *TTLL12* and *ZNF682* likely act at least in part via changes in immune cell composition.

### Immunogenetic response to parasite involves erythrocyte-related genes

We next tested whether red colobus genes with expression significantly correlated to parasitemia were united in having certain biological characteristics more likely than expected by chance. Using known orthologous human-colobus genes, we tested whether genes up-regulated in response to parasitemia were enriched for involvement in disease phenotypes (using Human Phenotype Ontology data; Köhler et al., 2019). The phenotypes with the strongest evidence of enrichment among our up-regulated genes included many red blood cell (RBC) disorders (Fig. 6; Table S8). These included stomatocytosis, or the production of RBCs with an abnormally shaped zone of central pallor (HP:0004446; adj. *p* = 0.00352); poikilocytosis (HP:0004447; adj. *p* = 0.0132), or the production of abnormally shaped RBCs; and reticulocytosis, or the increased production of immature erythrocytes which is disrupted during malaria anaemia in humans (HP:0001923; adj. *p* = 0.0257).

**Figure 6.**
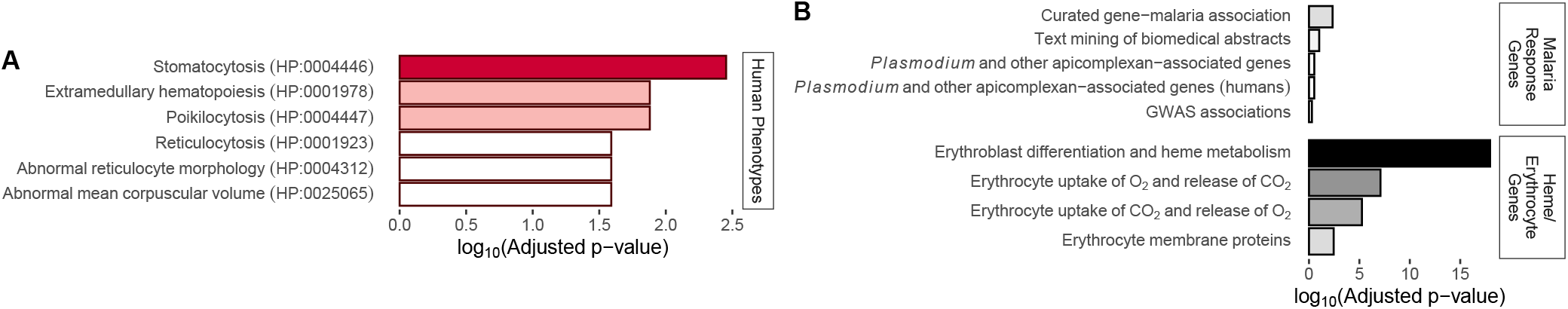
Overlap of colobus *Hepatocystis* response genes with human phenotype-associated genes and genes implicated in malaria response. Shading indicates strength of enrichment of Hepatocystis response genes with each functional category. **(A)** Over-represented human phenotypes, typically red blood cell disorders, were the result of a blind test for functional enrichment across all human phenotype annotation categories. **(B)** Malaria response gene sets were included *a priori* to compare *Hepatocystis* response to prior findings on *Plasmodium* response, while selection of heme and erythrocyte gene sets was informed by blind functional enrichment results.

Red colobus genes involved in *Hepatocystis* response were often implicated in all three of the enriched RBC disorders (Fig. S6). Such genes with significant association with *Hepatocystis* parasitemia (FDR adjusted *p <* 0.2) included *RHAG* (adj. *p* = 0.108), *SPTA1* (adj. *p* = 0.108), *SLC4A1* (adj. *p* = 0.137), *ABCB6* (adj. *p* = 0.163), and *SLC2A1* (adj. *p* = 0.174). Genes associated with a subset of the RBC phenotypes include *KLF1*, associated with reticulocyte disorders (adj. *p* = 0.122), as well as *STEAP3* (adj. *p* = 0.189) and *JAK2* (adj. *p* = 0.195), both linked to poikilocytosis. Many of these RBC disorder genes have been associated with malaria response in mice or humans. For instance, in mice vaccinated with erythrocyte ghosts isolated from *P. chaubaudi* -infected erythrocytes, *RHAG*, *SPTA1*, *KLF1*, and *SLC4A1* were among the erythropoiesis-involved genes that were significantly upregulated in response to infection with blood stage *Plasmodium chabaudi* (Delic et al., 2020). Furthermore, variation in *SPTA1*, which encodes a component of the RBC plasma membrane, was recently found to impact *P. falciparum* growth in cultured human RBCs (Ebel et al., 2020). Lastly, heterozygous deletion in *SLC4A1* causes South Asian hereditary ovalocytosis, which confers protection in humans against many *Plasmodium* species (Jarolim et al., 1991; Paquette et al., 2015), and this gene has been found to be under differential selection pressure in other primate species (Steiper et al., 2012).

In addition to the above blood-related disorders, the monkey genes with expression most strongly positively correlated to *Hepatocystis* parasitemia were also more likely than expected to be related to extramedullary hematopoiesis (HP:0001978; adj. *p* = 0.0132), or the production of cellular blood components outside of bone marrow.

Extramedullary or stress hematopoiesis is typically an indication of pathology in adults (Kim, 2010), and an increase in splenic erythropoiesis has been observed in mice with malarial anemia (reviewed in Lamikanra et al., 2007) and suggested as a mechanism by which humans compensate for anemia during *P. vivax* infection (del Portillo et al., 2004).

No RBC-related disorders were enriched among genes negatively correlated to parasitemia. No Gene Ontology (Ashburner et al., 2000) functional annotations were over-represented among either significantly up- or down-regulated genes in response to inferred *Hepatocystis* parasitemia (Table S8).

### Genetic architecture of *Hepatocystis* response overlaps that of *Plasmodium*

As the blood stage of *Hepatocystis* differs from that of *Plasmodium* in being limited to the gametocyte sexual stages, the *Hepatocystis* erythrocytic phase is predicted to not induce a cytokine response in the host (Ejotre et al., 2020). Despite this key difference, we found significant overlap between the genetic architecture of the colobus *Hepatocystis* response and the human response to malaria (Fig. 6; Table S9). Genes with significant increased gene expression response of colobus to *Hepatocystis* were significantly more likely than expected by chance to be associated with human response to malaria through curated gene-disease association (Davis et al., 2009; *p* = 0.00455) or text mining of biomedical abstracts, albeit not significantly (Pletscher-Frankild et al., 2015; *p* = 0.0963). However, *Hepatocystis* response genes did not significantly overlap malaria response genes defined via GWAS associations (Becker et al., 2004; *p* = 0.509), possibly highlighting the difficulty in generalizing GWAS results across species or populations. Down-regulated *Hepatocystis* response genes were enriched for only malaria response genes defined via text mining of biomedical abstracts (Pletscher-Frankild et al., 2015; *p* = 0.0128).

When intersected with genes implicated in mammalian response to *Plasmodium* and other apicomplexan parasites (Ebel et al., 2017; Fig. 6; Table S9), we found the overlap with upregulated colobus *Hepatocystis*-response genes to be no larger than expected by chance (*p* = 0.295). The concordance was no higher when the former was restricted to genes with evidence from human studies (*p* = 0.300), in contrast to our above results. The same was true for significantly down-regulated genes (*p* = 0.395 and *p* = 0.240 respectively.)

Prompted by the preponderance of RBC disorders among the enriched phenotypes, we next tested whether upregulated host genes were significantly enriched for association with selected erythrocyte functions (Fig. 6; Table S9). Significant enrichments included genes involved in erythroblast differentiation and metabolism of heme (*p* = 1.09 × 10^*−*18^), genes involved in the take up of oxygen and release of carbon dioxide (*p* = 8.11 × 10^*−*8^) and the reverse (*p* = 5.59 × 10^*−*6^), and genes encoding erythrocyte membrane protein genes (*p* = 3.57 × 10^*−*3^). Colobus genes that were down-regulated in response to *Hepatocystis* were not significantly enriched for these erythrocyte-related functional annotations (*p* = 1 for all). Although genes involved in interactions with red blood cells have been lost in *Hepatocystis* (Aunin et al., 2020), *Hepatocystis* has been reported to induce anemia in primates (Garnham, 1966; Chang et al., 2011), which may mediate the link between parasitemia and these monkey RBC-related genes.

### *Hepatocystis* response genes are selected in primates

Apicomplexan parasites such as malaria have exerted arguably the strongest selective force on humans (Kwiatkowski, 2005; Kariuki and Williams, 2020) and other primates (Tung et al., 2009; Oliveira et al., 2012; Ebel et al., 2017). The evolutionary arms race between host and pathogen or parasite may result in rapid evolution of gene sequences, and scans for signatures of natural selection in human and non-human genomes have repeatedly indicated immune genes to be the targets of adaptive evolution (Deschamps et al., 2016; Sironi et al., 2015; Fumagalli et al., 2011; Enard et al., 2016; Voight et al., 2006; Nielsen et al., 2005). In light of this, we tested whether red colobus genes involved in *Hepatocystis* response had accelerated evolution in primates (van der Lee et al., 2017). Colobus genes with expression that was positively correlated with *Hepatocystis* parasitemia were significantly more likely than expected by chance to be under positive selection in primates (*p* = 0.00847), but the same was not true for down-regulated genes (*p* = 1). Our findings are suggestive of potential *Hepatocystis*-mediated selection or generalized selection pressure from apicomplexan or haemosporidian parasites in primates.

### Few parasite genes have expression that varies with parasitemia

In parallel to our tests for colobus host genes that were associated with parasitemia, we also tested whether any *Hepatocystis* gene expression was similarly correlated with parasite load (Fig. S7; Table S10). The three genes with the strongest evidence had expression inversely proportional to parasitemia. Two were proteins with unknown function (HEP 00496100: *p* = 5.00 × 10^*−*5^, adj. *p* = 0.222; and HEP 00472500: *p* = 1.40 × 10^*−*4^, adj. *p* = 0.275), while the third (HEP 00452800: *p* = 1.86 × 10^*−*4^; adj. *p* = 0.275) is a putative circumsporozoite- and thrombospondin-related anonymous protein (TRAP)-related protein (CTRP), based on it exhibiting a thrombospondin type 1 domain (Pfam:PF00090.19). CTRPs have been previously implicated in parasite motility and host cell invasion, making them promising drug targets (Morahan et al., 2009). Our finding that this CTRP is most active when parasitemia is low may be linked to findings that TRAP is essential for early (liver) stage infection but not necessary for later (blood) stage infection in *Plasmodium* (Sultan et al., 1997).

### Parasitemia-associated co-expressed genes associated with lipid biosynthesis

Finally, to explore the interaction between colobus monkeys and *Hepatocystis* parasites from a systems biology perspective, we identified co-expressed colobus gene modules, and tested whether the expression of the modules was correlated with inferred *Hepatocystis* parasitemia or the expression of specific *Hepatocystis* genes. We identified two colobus gene modules for which the eigengene expression (expression profile summary) values were significantly correlated with inferred parasitemia (*p <* 0.01), one positively and one negatively (Table S11).

We found one cluster of 143 co-expressed colobus genes to be positively correlated with inferred parasitemia (*p* = 0.00277, *r*^2^ = 0.535), and these genes were enriched (Table S12) for being in the extracellular region (GO:0005576, *p* = 3.68 × 10^*−*5^) and for involvement in antimicrobial humoral immune response mediated by antimicrobial peptides (GO:0061844, *p* = 5.41 × 10^*−*3^; Table S12). No *Hepatocystis* gene had expression that was significantly correlated with the eigengene expression of this colobus module.

Another cluster of 215 genes had expression negatively correlated with inferred parasitemia (*p* = 0.00528, *r*^2^ = −0.504), and these genes were enriched for being involved in various processes, including biosynthesis of neutral lipids (GO:0046460, *p* = 5.20 × 10^*−*3^) specifically acylglycerol (GO:0046463, *p* = 5.20 × 10^*−*3^; Table S12). Acylglycerols (triacylglycerol and, secondarily, diacylglycerol) have been found to accumulate in erythrocytes infected with *P. falciparum* during the late trophozoite and schizont stages (Nawabi et al., 2003). Our findings are consistent with knowledge that *Plasmodium* parasites often hijack host lipid receptors, to invade other tissues, increasing infection severity (reviewed in Alsultan et al., 2020; Orikiiriza et al., 2017; O’Neal et al., 2020) and a recent study of malaria in Rwandan children which found a metabolic shift to increased production of long-chain unsaturated fatty acid associated triacylglycerols in response to infection with *P. falciparum* (Orikiiriza et al., 2017). The genes in this module with expression that most strongly correlated to parasitemia included *NAGA* (logFC = −4.63, *p* = 2.18 × 10^*−*6^, adj. *p* = 0.0133) and *TBC1D9B* (logFC = −1.33, *p* = 4.36 × 10^*−*5^; adj. *p* = 0.0454) which have been linked in humans to platelet count or count (via association of enhancers rs972578 or rs8137128; Astle et al., 2016) and blood protein measurement (via association of enhancer rs73351608; Sun et al., 2018), respectively. The expression of a large number of *Hepatocystis* genes (*N* =512) was significantly (*p <* 0.01) correlated with the eigengene expression of this module (Table S13), and these co-expressed *Hepatocystis* genes were enriched for protein channel activity (GO:0015252; *p* = 0.0476; Table S14).

### Conclusions and future directions

Overall, our study reveals considerable overlap between the colobus response to *Hepatocystis* and human response to *Plasmodium*. By studying the gene expression response to these parasites in non-human hosts, particularly our close non-human primate relatives, similar future studies will improve our understanding of broad patterns of host-parasite co-evolution. Such comparative context is critical for understanding the evolution of this important disease, and mechanistic insight into disease resistance may inform prioritization of possible intervention targets for human malaria patients. Understanding the genetic architecture of malaria response in a comparative context has implications for the determination of host specificity. This has additional practical importance, as multiple instances of non-human primate to human (*i.e.*, zoonotic) transmission have occurred. These zoonoses include, most pressingly, *Plasmodium knowlesi*, a monkey-origin malaria species which crossed over to humans in the late 20th century. We predict that similar future studies in diverse primate species that have co-evolved with these important parasites will further improve our understanding of host-parasite co-evolution.

## Materials and Methods

### Dataset

We downloaded RNA sequence (mRNA-seq) data previously generated (by Simons et al., 2019) from whole blood samples of 29 red colobus monkey (*Piliocolobus tephrosceles*) individuals (study PRJNA413051) from the National Center for Biotechnology Information (NCBI) Sequence Read Archive (SRA). Methods for sample collection, RNA extraction, library preparation, and transcriptome sequencing are described in Simons et al., 2019. Briefly, Simons and colleagues (2019) prepared mRNA-seq libraries with the KAPA Biosystems Stranded mRNA-seq kit and sequenced these on the Illumina HiSeq 4000 platform with 150 paired-end cycles.

### Gene expression inference

We removed Illumina-specific adapter regions (using parameters ILLUMINACLIP:TruSeq3-PE.fa:2:30:10) and low quality bases (quality score *<* 3) at the beginning and end of each raw sequencing read, and trimmed the read when the average quality of bases in a sliding 4-base wide window fell below 15 using Trimmomatic (Bolger et al., 2014). We removed reads that were less than 36 bp after trimming. In order to determine if sequence reads were from the colobus (GCF 002776525.3 ASM277652v3; Simons et al., 2019) or *Hepatocystis* (GCA 902459845.2 HEP1; Aunin et al., 2020), we performed a competitive mapping analysis as follows. Using the STAR alignment program (Dobin et al., 2013), we mapped trimmed, quality-filtered RNA-seq sequencing reads to concatenated reference genomes that include colobus and *Hepatocystis* genomes, allowing each read to map multiple locations with up to eight mismatches per 100 bp paired-end read (after Lee et al., 2018). The best matching, primary alignments were extracted and then merged into pathogen and host “unique” datasets (*i.e.*, primary reads that have only mapped to the host or parasite), while excluding unplaced scaffolds for colobus. Libraries which had been split and sequenced in multiple lanes were then combined by red colobus monkey individual (*N* =29).

We used the Rsubread R package (Liao et al., 2019) to create a colobus monkey read-count matrix for each sample, using the “uniquely” mapped host reads and allowing for overlapping reads. Reads were tallied at the level of exons which were then summed to create a per-gene count for the colobus monkey, using gene annotations (GCF 002776525.3 ASM277652v3). For *Hepatocystis*, we used published read counts per gene to be concordant with prior work (Aunin et al., 2020). Read counts per gene and the results of downstream analyses were largely consistent between analyses using the published *Hepatocystis* read count dataset and sums of read counts per gene from our mapping analysis (supplemental text).

### Inference of parasite load

We inferred a proxy of parasitemia by calculating the ratio of the number of pathogen reads (data from Aunin et al., 2020) to the number of host colobus reads recovered (after Yamagishi et al., 2014; Table S1; Fig. 1). For this measure, we used the *Hepatocystis* read counts for genes, excluding the upper and lower quartiles of the distribution to avoid noise from genes with extremely high or low expression. However, using the full set of genes yielded very similar results (supplemental text).

### Tests for cell type or stage correlates to inferred parasitemia

Using our inferred *Hepatocystis* parasitemia and previously published relative immune cell composition abundance data (Simons et al., 2019), we tested whether parasitemia and immune cell proportions were correlated by testing the hypothesis that Pearson’s correlation coefficient is not equal to zero using a *t*-test. We applied the same approach to test for association between inferred parasitemia and fraction of *Hepatocystis* cells in each developmental stage, which we estimated via bulk RNAseq deconvolution as follows.

In order to infer the ratio of different *Hepatocystis* cell types (*i.e.*, life stages) in our bulk data and determine how these may covary with parasitemia and the expression of differentially expressed genes, we deconvoluted the bulk *Hepatocystis* RNAseq read count data. To do so, we used published stage-specific “pseudo-bulk” samples (Aunin et al., 2020) constructed from data gathered from the Malaria Cell Atlas (Howick et al., 2019), which was based on single cell sequencing of cells in known life stages to deconvolute previously published *Hepatocystis* read count data. For the analysis, we used relationships between *Hepatocystis* genes and their one-to-one orthologous *Plasmodium berghei* genes, which were determined by Aunin et al., 2020 using OrthoMCL (Li et al., 2003) and IQ-TREE (Nguyen et al., 2014). We excluded those *Hepatocystis* genes without one-to-one orthology with *P. berghei* genes. All analyses for estimating cell type proportions in our *Hepatocystis* read count data were implemented in BSeq-sc (Baron et al., 2016) and CIBERSORTx (Newman et al., 2019) which we ran with 1,000 permutations (Table S3). For downstream analyses, we combined all non-erythrocytic life stages into an “Other” category to remain consistent with previously published data.

Finally, we tested for an association between these relative abundances of *Hepatocystis* developmental stages inferred via bulk RNAseq deconvolution and colobus immune cell types (Simons et al., 2019) using a *t*-test to establish if Pearson’s correlation coefficient differed significantly from zero.

### Identification of differentially expressed genes by parasitemia

To identify host colobus monkey and *Hepatocystis* genes with expression correlated to parasitemia, we analyzed the two datasets independently with the Bioconductor package edgeR (Robinson et al., 2010) as follows. We first excluded low coverage genes, defined as any gene with expression below a counts per million (CPM) threshold corresponding to a read count of 10 in at least two individuals for *Hepatocystis* genes or corresponding to a read count of 5 in at least four individuals for colobus genes. 12,223 colobus genes and 4,432 *Hepatocystis* genes remained after filtering. We then normalized the read count matrices to control for library size and compositional bias.

From the read-count matrices for the host and parasite separately, we identified significantly differentially expressed genes with expression correlated with parasitemia (“parasitemia-DE genes”) using the edgeR package (Robinson et al., 2010) as follows. After estimating dispersion using the Cox-Reid profile-adjusted likelihood (CR) method and fitting a negative binomial model, we identified significantly correlated genes using the generalized linear model (GLM) likelihood ratio test. To do so, we used the model:

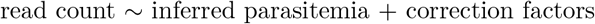

where correction factors include those adjusting for differences in number of reads and library complexity. We corrected for multiple tests via false discovery rate (FDR) estimation. We considered a gene to be a parasitemia-DE gene if the adjusted *p*-value was less than 0.2 for colobus genes, or if the unadjusted *p*-value was less than 0.01 for *Hepatocystis*, using a lower threshold due to reduced statistical power in analyses of parasite gene expression.

To test for an influence of immune cell composition on differential expression by parasitemia, we repeated the above analysis adding in parameters for relative abundance of memory B cells, CD8 naïve CD4+ T cells, memory resting CD4+ T Cells, memory activated CD4+ T cells, natural killer cells, monocytes, macrophages, and neutrophils. All fractions of colobus immune cells were taken from Simons et al., 2019 and are based on bulk RNAseq deconvolution analysis. The filtering and testing for differential expression by parasitemia took place as in the simpler model without cell type composition parameters. Results are presented joined with those of the simpler differential expression analysis (Table S7).

### Functional enrichment analyses

We next tested whether the genes with expression that was significantly positively correlated with parasitemia (upregulated parasitemia-DE genes) or significantly negatively correlated with parasitemia (downregulated parasitemia-DE genes) were enriched for particular biological functions. Specifically, we tested whether these genes were more likely than expected by chance to share Gene Ontology molecular functions, biological processes, or cell component annotations (Ashburner et al., 2000; The Gene Ontology Consortium, 2019). For the colobus dataset, we also tested for functional enrichment of Human Phenotype Ontology (Köhler et al., 2019) annotations, using annotations for human orthologs matched to our colobus genes using the Ensembl gene mappings (accessed via g:Orth; Raudvere et al., 2019). To test for enrichment, we used a hypergeometric test as implemented in g:Profiler (Raudvere et al., 2019) and used all annotated genes after filtration for low read count as a comparative background list for differential expression determination.

To compare colobus response to *Hepatocystis* with parasite response in humans and other mammals, we further analyzed the colobus dataset as follows. We tested whether there was significantly more overlap than expected by change between our up- or downregulated parasitemia-DE genes and particular sets of malaria- or blood-related genes. As these genes sets were based on human gene identifiers, we used all human genes that had orthology with colobus genes sets for our significantly up- and down-regulated genes, as well as with a comparative background set all colobus genes with read count data after filtering. The sets of *a priori* genes related to malaria included genes from curated gene-malaria association (*N* =19; Davis et al., 2009), genes associated with malaria through text mining of biomedical abstracts (*N* =701; Pletscher-Frankild et al., 2015), and malaria response genes as defined via GWAS associations (*N* =51; Becker et al., 2004). We also tested sets of genes implicated in mammalian response to *Plasmodium* and related parasites (*N* =490; Ebel et al., 2017), and the same dataset reduced to only include those with evidence of association in humans (*N* =301). Our final gene sets were selected due to their connections with erythrocyte functions. Gene sets were sourced from MSigDB (Subramanian et al., 2005; Liberzon et al., 2015) and included genes involved in erythroblast differentiation and metabolism of heme (*N* =200; MSigDB ID M5945), genes involved in the uptake of oxygen and release of carbon dioxide (*N* =11; MSigDB ID M29536; Jassal et al., 2020) and the uptake of carbon dioxide and release of oxygen (*N* =16; MSigDB ID M26928; Jassal et al., 2020), and genes encoding erythrocyte membrane protein genes (*N* =14; MSigDB ID M2442; Steiner et al., 2009).

### Evolutionary analyses

Finally we tested whether the colobus parasitemia-linked genes were enriched for genes previously inferred to be under positive natural selection in primates. As above, we tested for greater overlap than expected by chance between our up- and downregulated colobus parasitemia-DE genes and a list of 331 protein-coding genes previously found to be under strong positive selection (*i.e.*, having a significantly better fit for a selection model than a neutral model in a maximum likelihood d*N* /d*S* analysis and containing at least one significant positively selected amino acid residue) in a comparative analysis of nine whole-genome sequenced primates (van der Lee et al., 2017).

### Identification of co-expressed gene modules associated with parasitemia

To identify key sets of genes involved in host-parasite interactions between colobus and *Hepatocystis*, we inferred gene co-expression networks using Weighted Gene Co-expression Network Analysis of the colobus genes (WGCNA; Langfelder and Horvath, 2008). We used the same read count matrix for the identification of parasitemia-DE genes. For clustering, we selected a soft-thresholding power (*β*) via analysis of network topology, visually examining scale-free topology fit index as a function of soft-thresholding power and selecting *β* = 9 as the value for which the scale free topology model fit *R*^2^ is approximately 0.9 (*R*^2^ = 0.894).

We performed automatic network construction and module detection in a block-wise manner using the WGCNA function blockwiseModules. Signed network options were selected for the adjacency function (networkType = “unsigned”) and topological overlap (TOMType = “signed”). For tree cutting, the minimum module size was set to 30, and for module merging, the merge cut height was set to 0.15. The gene reassignment threshold was set to 0.

Using the same parasitemia proxy as the differential expression analysis, we next tested for co-expression modules with expression correlated to parasite load by calculating the Pearson correlation coefficient and computing a *p*-value via the Student’s *t*-test with asymptotic null distribution. For each significant module, we identified significantly over-represented functional annotations using a hypergeometric test as implemented in g:Profiler (Raudvere et al., 2019), using the same parameters as the previous function enrichment analysis of parasite-DE colobus genes. Similarly, we also used all annotated genes after filtration for low read count as a comparative background list.

Finally, we tested if any significantly parasitemia-associated modules co-expressed with *Hepatocystis* gene expression using the same read count data as the differential expression analysis above. To do so, we tested for a correlation between the colobus module’s eigen expression and *Hepatocystis* gene expression using Pearson’s product moment correlation coefficient and testing for significance using a *t*-test. For each module, we next tested whether *Hepatocystis* genes with correlated expression were enriched for biological annotations. To associate genes with functional annotations, we replaced *Hepatocystis* gene identifiers to their one-to-one *Plasmodium berghei* orthologues, as inferred by Aunin et al. (2020) using OrthoMCL (Li et al., 2003). We identified significantly over-represented functional annotations using a hypergeometric test as implemented in g:Profiler (Raudvere et al., 2019), using all annotated genes as a comparative background list.

## Code availability

RNAseq data have been previously published (Aunin et al., 2020) and released via the NCBI Sequence Read Archive under accession PRJNA413051. Supplemental tables, figures, and text are published under a Creative Commons Attribution 4.0 International license and available online (doi:10.5281/zenodo.4287878). Code for all analyses is available in a repository (doi:10.5281/zenodo.4287572; https://github.com/ambertrujillo/Colobus_dualRNAseq) and released under the open source GNU General Public License v3.0.

## Acknowledgements

This work was supported by the Ford Predoctoral Fellowship to A.E.T. We thank Alex DeCasien for assistance with bulk RNA deconvolution analyses. This work was supported in part through the NYU IT High Performance Computing resources, services, and staff. The authors acknowledge the Office of Advanced Research Computing (OARC) at Rutgers, The State University of New Jersey for providing access to the Amarel cluster and associated research computing resources that have contributed to the results reported here.

## Supplemental Figure and Table Captions

### Supplemental Table Captions

**Table S1. Inferred parasitemia by sample.** Competitively mapped, unique colobus read counts and previously published *Hepatocystis* read counts from each colobus sample.

**Table S2. Immune cell type proportion correlation with parasitemia.**

Pearson coefficient for correlation between colobus immune cell type proportions and inferred *Hepatocystis* parasitemia, along with *t*-test for significant association (deviation from zero). Proportions of immune cell types are from Simons et al., 2019 and include memory B cells (“MemB”), CD8 Naïve CD4+ T Cells (“naiveCD4plus”), Memory Resting CD4+ T Cells (“MemrestCD4plus”), Memory Activated CD4+ T Cells (“MemactCD4plus”), natural killer cells (“restNK”), monocytes, macrophages, and neutrophils.

**Table S3. Inferred *Hepatocystis* life stage proportions for each sample.**

Inferred *Hepatocystis* life stage proportions for each colobus sample after deconvolution analysis using a cell atlas based on *Plasmodium berghei* single-cell data.

**Table S4. *Hepatocystis* life stage proportion correlation with parasitemia.** Pearson coefficient for correlation between *Hepatocystis* developmental stage proportions and inferred *Hepatocystis* parasitemia, along with *t*-test for significant association (deviation from zero).

**Table S5. Results of test for correlation between *Hepatocystis* life stages and immune cell composition.** Pearson coefficient for correlation between *Hepatocystis* developmental stage proportions and colobus immune cell type proportions, along with *t*-test for significant association (deviation from zero). Proportions of immune cell types are from Simons et al., 2019 and include memory B cells (“MemB”), CD8 Naïve CD4+ T Cells (“naiveCD4plus”), Memory Resting CD4+ T Cells (“MemrestCD4plus”), Memory Activated CD4+ T Cells (“MemactCD4plus”), natural killer cells (“restNK”), monocytes, macrophages, and neutrophils.

**Table S6. Colobus genes differentially expressed by parasitemia.** Full results of test for differential expression of colobus genes in response to *Hepatocystis* parasitemia. The log_2_ fold change in response to inferred parasitemia (“logFC”) is computed per unit change in inferred parasitemia proxy.

**Table S7. Colobus genes differentially expressed by parasitemia controlling for colobus immune cell type proportions.**

**Table S8. Functional enrichment of colobus genes DE by parasitemia.**

**Table S9. Comparison of colobus *Hepatocystis*-response genes with malaria response-associated gene sets.** Significantly up-regulated *Hepatocystis* response genes are overrepresented in some sets of genes connected to *Plasmodium* malaria as well as those associated with erythrocytes.

**Table S10. *Hepatocystis* genes differentially expressed by parasitemia.** Full results of test for differential expression of *Hepatocystis* genes in response to *Hepatocystis* parasitemia. The log_2_ fold change in response to inferred parasitemia (“logFC”) is computed per unit change in inferred parasitemia proxy.

**Table S11. Co-expression modules correlated with parasitemia.** Clusters of genes with similar co-expression, inferred via weighted gene co-expression network analysis (WGCNA). Correlation with parasitemia summarized by *p*-value indicating result of *t*-test that Pearson’s correlation coefficient is not equal to zero.

**Table S12. Co-expression module functional enrichment.** Functional categories (Gene Ontology and Human Phenotype) overrepresented among colobus genes in the two co-expression modules with the strongest evidence for correlation with parasitemia.

**Table S13. *Hepatocystis* genes with expression correlated to that of colobus co-expression modules.** Correlation with parasitemia summarized by *p*-value indicating result of *t*-test that Pearson’s correlation coefficient is not equal to zero.

**Table S14. Functional enrichment of *Hepatocystis* genes with expression correlated to that of colobus co-expression modules.** Functional categories (Gene Ontology) overrepresented among *Hepatocystis* genes that contrived with the colobus co-expression modules with the strongest evidence for correlation with parasitemia.

### Supplemental Figure Captions

**Figure S1. Relationship between relative abundance of immune cell types and inferred *Hepatocystis* parasitemia.** Proportions of immune cell types are from Simons et al., 2019 and include memory B cells (“MemB”), CD8 naïve CD4+ T cells (“naiveCD4plus”), memory resting CD4+ T cells (“MemrestCD4plus”), memory activated CD4+ T cells (“MemactCD4plus”), natural killer cells (“restNK”), monocytes, macrophages, and neutrophils.

**Figure S2. Proportion of *Hepatocystis* reads inferred to be in each parasite developmental stage via deconvolution.** Deconvolution based on previously published pseudobulk single-cell *Plasmodium berghei* data constructed from the Malaria Cell Atlas. Results are grouped by colobus individual blood sample. Proportions are consistent with previously published data.

**Figure S3. Relationship between inferred parasitemia and parasite life stage proportion.** Proportions shown for schizonts, merozoites, ring-form trophozoites, mature trophozoites, female and male gametocytes, and other.

**Figure S4. Expression of colobus *ACKR1* gene plotted against inferred *Hepatocystis* parasitemia.** Expression summarized as normalized read count per million reads sequenced. One outlier data point was removed from the plot.

**Figure S5. Comparison of results of test for DE genes using model with and without immune cell composition parameters.** Points shown indicate unadjusted *p*-value for each test, plotted on a negative logarithmic scale. Red points were significant in the simple model without immune cell composition parameters (FDR-adjusted *p <* 0.05). Only *ANP32E* and *SLC18A2* (labeled) were significant at this strict threshold (FDR-adjusted *p <* 0.05) in the more complex model that included cell composition parameters.

**Figure S6. Overlap of parasitemia-DE genes associated with RBC disorders.** Various red blood cell-related disorders had the strongest evidence of enrichment among parasitemia-DE genes. Heatmap shows overlap among their associated up-regulated genes. The degree of log-fold change is indicated by the color gradient, while grey squares indicate no association between gene and phenotype.

**Figure S7. Volcano plot showing magnitude and significance of *Hepatocystis* gene expression changes correlated with inferred *Hepatocystis* parasitemia.** Labelled red circles indicate *Hepatocystis* genes significantly up- or down-regulated in response to parasitemia (uncorrected *p <* 0.05). The log_2_ fold change in response to inferred parasitemia (x-axis) is computed per unit change in inferred parasitemia proxy.

